# Predicting Adverse Drug-Drug Interactions with Neural Embedding of Semantic Predications

**DOI:** 10.1101/752022

**Authors:** Hannah A. Burkhardt, Devika Subramanian, Justin Mower, Trevor Cohen

## Abstract

The identification of drug-drug interactions (DDIs) is important for patient safety; yet, compared to other pharmacovigilance work, a limited amount of research has been conducted in this space. Recent work has successfully applied a method of deriving distributed vector representations from structured biomedical knowledge, known as Embedding of Semantic Predications (ESP), to the problem of predicting individual drug side effects. In the current paper we extend this work by applying ESP to the problem of predicting polypharmacy side-effects for particular drug combinations, building on a recent reconceptualization of this problem as a network of drug nodes connected by side effect edges. We evaluate ESP embeddings derived from the resulting graph on a side-effect prediction task against a previously reported graph convolutional neural network approach, using the same data and evaluation methods. We demonstrate that ESP models perform better, while being faster to train, more re-usable, and significantly simpler.

## Introduction

Drug-drug interactions (DDIs) are an important challenge to patient safety. Patients are commonly prescribed multiple drugs at a time, a phenomenon known as polypharmacy. The incidence of polypharmacy has been increasing^1^, and is generally very high in some patient groups, such as the elderly – the CDC reported that between 2011 and 2014, 66.8% of those 65 years and older take three or more prescription medications^2^. While multiple drugs can enhance each other’s efficacy and may be given in combination intentionally, many drug combinations can have unintended and harmful physiological consequences^3^. For example, if two medications affect the same biological pathway, they might compete for a substrate and the effect of one or both might be decreased or altered, side effects might be exacerbated, or new side effects might result from the drug interaction.

Identifying and accounting for polypharmacy effects is difficult^1,3^. While new drugs are tested for side effects during clinical trials, it is not feasible to investigate every possible combination of the new drug with all drugs currently on the market, let alone all investigational agents. As clinical trials do not exhaustively capture even the side effects of individual drugs, pharmacovigilance systems have been put in place for post-market surveillance to monitor for, and support regulatory action against, harmful drug effects. One such system is the Food and Drug Administration’s Adverse Event Reporting System (FAERS)^4^. Physicians, pharmacists, and others submit Adverse Drug Event (ADE) reports, which can then be analyzed for patterns of reporting, including identification of putative polypharmacy effects^5^. It is challenging to analyze these reports manually, so a range of data mining methods have been developed to detect drug/ADE associations^6^. Of importance for the current work, Tatonetti et al. developed a comprehensive database of DDI side effects, called TWOSIDES, using a data mining approach leveraging adverse event reports^5^.

Because of the combinatorial explosion of possible drug pairings to investigate, the development of computational methods to predict new polypharmacy side effects is a key task in pharmacovigilance. Several approaches to predicting drug-drug interactions have been described. Some approaches harness the idea that similar drugs will take part in similar drug interactions^6^. For this purpose, modeling of chemical and structural similarities has been explored^7,8^. Other approaches employ unstructured text, such as the Medline biomedical literature corpus, to extract interaction information with text mining techniques^9^. A third group of approaches to the DDI prediction problem uses binary classification techniques in order to classify a given drug pair as interacting or not interacting. A promising way to do this is to consider DDI prediction a link prediction problem, where drugs are nodes and side effects are the links between them^6,10^. However, predicting not only the presence of adverse effects of DDIs, but also the nature of such side effects, has not been extensively explored. For a detailed account of previously reported DDI prediction methods, the interested reader is referred to a recent comprehensive review by Vilar, Friedman and Hripcsak^3^.

The current work is inspired by Zitnik et al.’s graph convolutional network approach called Decagon^11^. Decagon makes use of the conceptualization of drug interactions as a graph, where drugs are nodes and side effects are edges, with different side effects being represented by edges of different types. Identifying polypharmacy side effects then becomes a multi-relational link prediction task. Graph embeddings are learned for drugs (nodes) as well as side effects (edges), and then used to predict which types of links are likely to exist between pairs of drugs. It is important to note that not only the presence of an interaction is predicted, but also the nature of this interaction, a distinguishing feature from other approaches such as Fokoue et al.’s link prediction model^10^. In fact, Decagon is reportedly the first approach to modeling different polypharmacy side effect types^11^. Decagon is trained on data from TWOSIDES, as well as various information sources for drug target and protein-protein interaction data, and achieves high performance as measured by area under the receiver operating curve (AUROC) and area under the precision-recall curve (AUPRC) for the task of predicting side-effects given a drug pair. On this task, Decagon outperforms several baseline multi-relational link prediction approaches by at least 0.1 in mean AUROC across ADEs: RESCAL^12^, a multi-relational tensor factorization approach involving the decomposition of a matrix encoding drug-drug relationships into components representing drugs and side effects, achieves a mean AUROC of 0.693; and DeepWalk^13^, a procedure which produces neural embeddings based on a biased random walk through the node neighborhood within the network and then uses these embeddings as a basis for logistic regression, achieves a mean AUROC of 0.761. Additionally, Decagon is used to predict novel DDIs, a number of which were supported by the literature. However, this approach also has limitations. First, node embeddings are learned with a graph convolutional network^14^ approach in which d-dimensional node embedding vectors are linked by d^2^-dimensional edge matrices. The computational demands of training such graph convolutional networks is high, limiting applicability to larger datasets – a problem that will not be eliminated by increasing compute power, as data set size increases at a similar rate^15^. Second, while the process produces vector embeddings for drugs (nodes), side-effect (edge) embeddings are represented by matrices and are thus challenging to re-use as they exist in a space of different dimensionality to the resulting drug vector space. Third, a simpler model might have advantages over Decagon because of its parsimony: if a smaller number of parameters is sufficient to model the underlying data, models with fewer parameters may avoid overfitting, and generalize better.

In the current work we propose an alternative approach to DDI prediction which builds upon Zitnik et al.’s graph conceptualization of the DDI prediction problem, but uses Embedding of Semantic Predications (ESP) ^16^, which can address each of these challenges, for representation learning. ESP is a method for generating vector embeddings for biomedical concepts, such as drugs and side effects, from concept-relationship-concept triples called predications, using a neural network. ESP makes use of vector symbolic architectures, in which the composition (binding) and addition (superposition) operators are used to create transient embeddings for composite concepts (such as INHIBITS cytochrome p-450) from their component vectors. During training, embeddings are updated such that vectors for concepts occurring in predications with similar predicate-argument pairs become more similar (closer in vector space) to each other. Conversely, vectors for dissimilar concepts approach orthogonality in the vector space. The high dimensionality of vectors in ESP (e.g. 10,000 bits in binary vector implementations) makes it unlikely for vectors to be similar to each other by chance alone^17^ – an n-dimensional vector space contains 2^n^ nearly orthogonal vectors^18^. Vectors are randomly initialized before the encoding process begins. For each positive triple in the training set, updates are made to the weight vectors representing the subject (S(*s*)), predicate (P(*p*)), and object (C(*o*)) as follows:

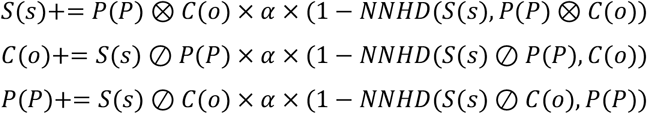

Note that, while ESP learns only one embedding for relationships (the *predicate vector P*), two embeddings are learned for each concept: the *semantic vector* (*S)* and the *context vector* (*C*). ⊗ is the binding operator, ⊘ is the release operator (inverse of binding), and + is the superposition operator. The implementation of these operators depends on the vector symbolic architecture approach used. Here we use the Binary Spatter Code^16,17^, in which binding is the elementwise XOR. Because XOR is its own inverse, the release and binding operators are equivalent in this implementation. The superposition operator, which allows multiple vectors to be “added” together, retains the elementwise most common element (1 or 0) across the superposed vectors, with ties broken at random. The update steps are also modulated by a linearly decreasing learning rate (*α*) and the similarity between the desired and current vectors, measured as the non-negative normalized Hamming Distance (NNHD)^16^. Analogous update steps are completed for negative samples, which are created by substituting random objects for the objects in positive triples (using -NNHD rather than 1-NNHD). The optimization objective, which is achieved through stochastic gradient descent, is the cross-entropy function defined as follows:

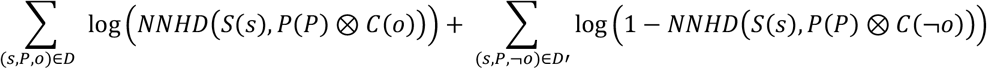

where (*s*, *P*, ¬*o*) is a training triple and (*s*, *P*, ¬*o*) is a training triple with corrupted object (a negative example). In words, the optimization objective is to create representations of concepts that minimize the distance between subjects and representations of their corresponding predicate-object pairs, while maximizing the distance between subjects and representations of predicate-object pairs that they do not correspond to. An illustration of the ESP architecture used to learn the embeddings is shown in Figure 1.

This network trains matrices of weights consisting of the vector embeddings for each drug, activated separately during training using one-hot vector representations. For example, when encoding warfarin-BLEEDING-aspirin, the input and output vectors are one-hot vectors, with a 1 in the warfarin position and a 1 in the aspirin position, respectively. When multiplying the vector by the weight matrix, all columns are set to 0, except the ones pertaining to these particular drugs. Consequently, only the weight matrix components for the concepts and predicate of a triple are involved in the update step that encodes it. This update proceeds toward the optimization objective shown above. The weight matrices are updated so as to move *S*(warfarin) closer to *P*(bleeding) ⊗ *C*(aspirin); *C*(aspirin) closer to *S*(warfarin) ⊘ *P*(bleeding); and *P*(bleeding) closer to *S*(warfarin) ⊘ *C*(aspirin). We refer the interested reader to Cohen et al.^16^ for a detailed exposition of the algorithm and evaluations across a range of pairwise entity-level tasks.

**Figure 1.**
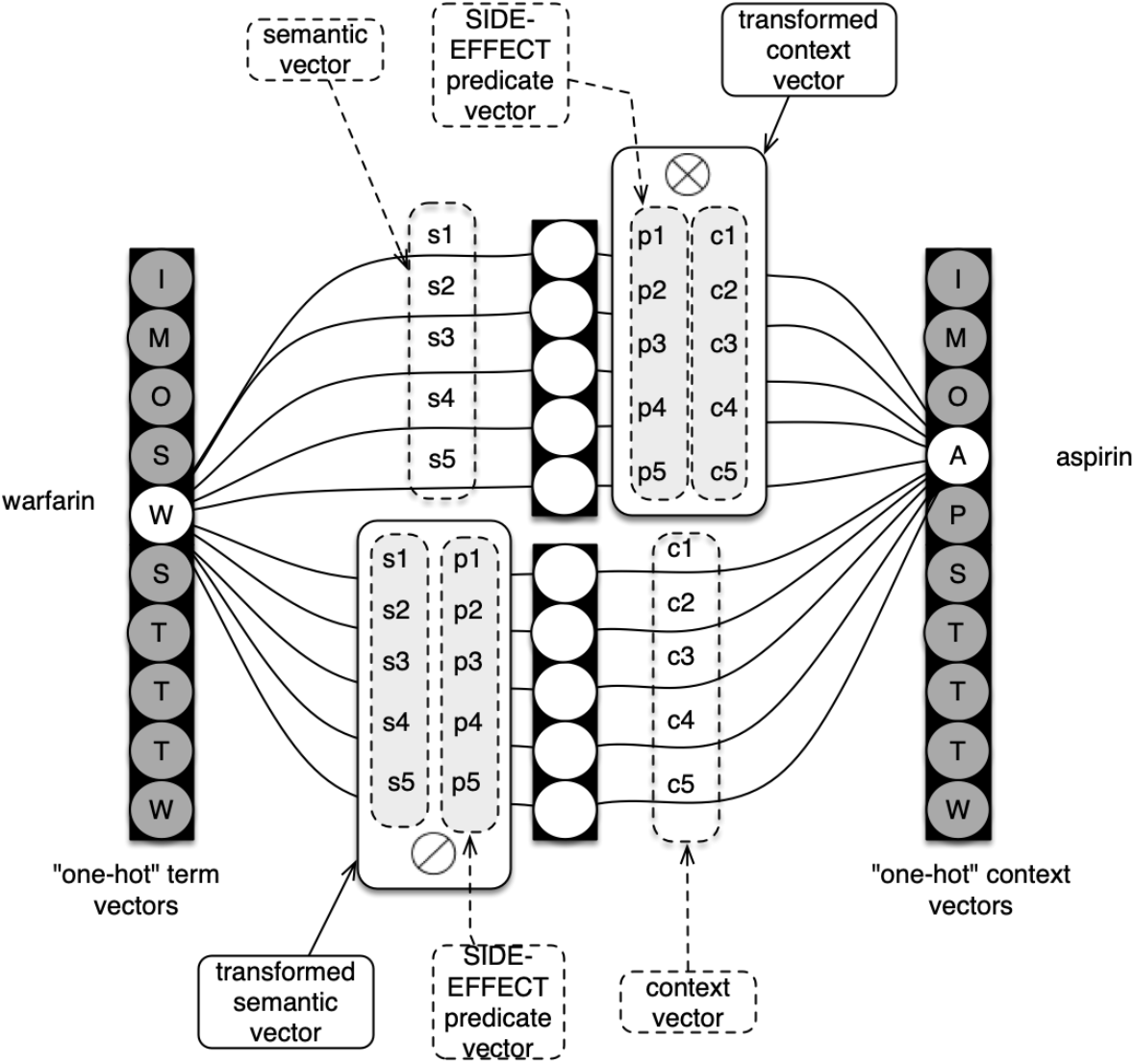
Illustration of the ESP architecture used to generate 5-dimensional embeddings for a 10-concept vocabulary. Adapted from Cohen et al.^16^

Because similar representations will be learned for similar concepts, we can query the resulting vector space for concepts related to (similar to) a given concept (or concept composition). In our approach, we generate representations for drugs and side effects based on drug/side-effect/drug triples, resulting in input weights which serve as semantic vectors and output weights which serve as context vectors for drugs, and predicate vectors for side effects. In the generated vector space, the bound product of the semantic and context vectors for two drugs should then be similar to the predicate vectors of the polypharmacy side effects that might be observed when the two drugs are taken in combination. A query of interest might thus be: “What polypharmacy side effects might be expected when taking warfarin and aspirin together?” The response is the predicate vector most similar to the bound product of S(warfarin) and C(aspirin) over all possible side effects, minimizing *NNHD*(*P*(side effect), *S*(warfarin) ⊗ *C*(aspirin)).

ESP has previously been used successfully in predicting side effects for individual drugs^16^. Most recently, Mower et al.^19^ utilized concept-relationship-concept triples from SemMedDB, a knowledge base extracted from the MEDLINE corpus of medical literature using SemRep^20^, to construct an ESP model which provided input to a supervised machine learning model trained to predict which side effects might be seen for which drugs, achieving strong results with an area under the receiver operating curve (AUROC) of 0.96 and 0.95 across two different test sets. The success of ESP for predicting individual drug side effects raises the question of whether the approach might be used to predict DDIs as well. In this paper, we explore the hypothesis that ESP has utility for the polypharmacy side effect prediction task.

## Methods

When selecting data for pharmacovigilance research, balancing level of curation with representation of emergent signals (such as those that might be found in ADE reports) is challenging^21^. As a result, prior evaluations of pharmacovigilance approaches have used a range of diverging data sets for training and evaluation. Because of the large variety of available data and their particular biases and limitations, it can be difficult to directly compare the performance of pharmacovigilance models^3^. To minimize this problem, we elected to use the same dataset used for evaluation of a recently published approach to identifying specific drug-drug adverse effects, to facilitate direct comparison of our results. Commendably, Zitnik et al. published both the data sets used in their approach and their code. As a result, we were able to use the same data as well as the same train/test splits, as produced by the publicly available code. The data were downloaded from Stanford’s SNAP website^22^ and the code was downloaded from Marinka Zitnik’s GitHub page^23^. We performed the additional preprocessing steps described by Zitnik et al., taking care to follow their methods as closely as possible. This included discarding any triples for side effects that occurred in fewer than 500 drug interaction triples, reducing the total number of side effects in the dataset from 1317 to 963.

After modifying the Decagon code to output the testing set as well as the data subset used for training, we allowed Decagon to complete several epochs of training. However, in our own execution of the protocol, we were not able to run Decagon for up to 100 epochs as described by Zitnik et al. (with documented early stopping) within a reasonable time frame, possibly due to differences between our groups’ computational resources. With a single NVIDIA Tesla p40 GPU we observed training times of roughly 36 hours per epoch, with a one-time up-front setup and initialization time of about 6 hours. Consequently, we obtained predictions over the testing set after allowing Decagon to train for 12 epochs only. We present both these results and the previously-published Decagon results showing best-documented performance, enabling fair comparison within constraints of our timeframe and computational resources.

The downloaded data set consists of 9,643,506 triples, of which 7,323,790 are drug-drug interaction triples, 2,289,960 are protein-protein interaction pairs, and 29,756 are drug target pairs (linking drugs to proteins). While the downloaded data set also contained individual drug side effects, these data are not used in the published code and were consequently left out for the purposes of the current research as well. The training data consist of 80% of the total data, with the remaining 20% being equally divided between the validation and test sets. In Decagon, the validation set is used to determine the early stopping point by computing the cross-entropy validation loss over the validation triples after each epoch; training is stopped early if validation loss does not improve for 2 consecutive epochs. Testing is performed exclusively over the testing set triples. It is of note that in the original Decagon work, the authors doubled the size of the dataset in the downloadable data files by also including all training triples in their inverted form, treating the “reverse” side effect edge types as separate edge types, resulting in a total of 963*2=1926 side effects (drug-drug edge types). Later, for purposes of analysis, we examine AUCs per side effect, only considering the original 963 side effects.

From these data, we created subject-predicate-object triples (predications) as input for ESP training (Table 1).

**Table 1.**
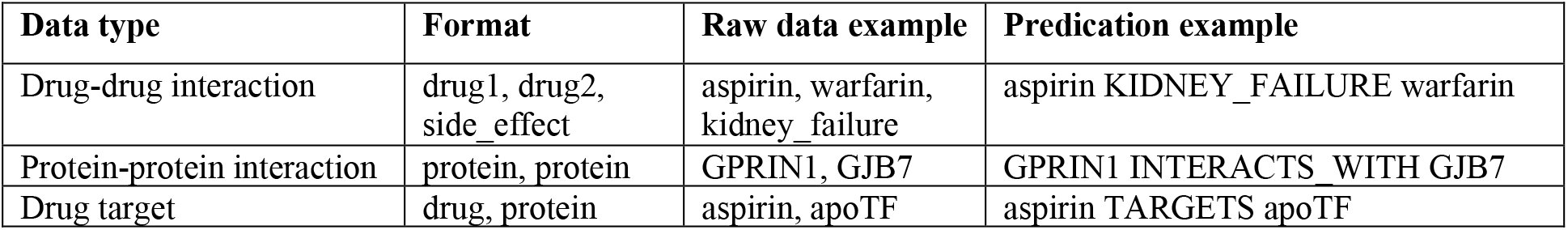
Predication examples. Vocabulary codes are replaced with names for the purpose of illustration.

An ESP model was trained using the open-source Semantic Vectors package^15,24,25^ pre-release version 5.9 with vector dimensionality of 16,000 bits. Unlike previous releases, this version allows predicate vectors to be trained (in addition to semantic and context vectors). This capability is essential for our approach to DDI prediction, as we generate trained embeddings for side effects (represented as predicates), as well as drugs.

We converted the testing data set to a list of similarity comparisons to be made. For each triple in the test set, we bind the semantic vector for the first drug with the context vector for the second drug and compare the similarity of the resulting compositional vector with the predicate vector for the side effect:

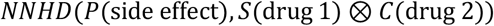

The result is a similarity score between 0 and 1. Identical vectors score 1 and orthogonal vectors 0. It is important to note here that the publicly available Decagon data sets, like most drug-drug interaction data sets, only contain positive examples, i.e. data points with a ground truth label of 1. ESP originates from methods that learn representations of words from free text, where the same problem exists; ESP therefore automatically performs negative sampling in the training process to account for this. However, during testing of ESP, we must test using generated negative examples. The published Decagon code generates negative examples by choosing 2 random drugs and connecting them with a random side effect, making sure that the resulting triple (edge) is not observed in the original, positive data. Additionally, for each side effect, we generate the same number of negative examples as we have positive examples. The negative testing examples produced by the Decagon code were used for evaluating the ESP model.

Evaluation was performed with Anaconda^26^ version 4.6.8 and Python version 3.6.8 using the scikit-learn package version 0.19.1. As in the original Decagon protocol, AUROC and AUPRC were calculated on a per side effect basis. Precision at *k* was also calculated, with *k*=50, and all three metrics were averaged across all side-effects.

While 16,000-bit vectors have been shown to perform well on a range of prior ESP evaluations, training time and model parsimony might be improved by using lower dimensional vectors. In order to determine the minimal space requirements of a model with acceptable accuracy, we analyzed several lower embedding dimensionalities. Due to the distributed nature of the vector representations (information is spread as a pattern across the entire vector), it is possible to test the loss in accuracy for smaller dimensional vectors without re-training, but rather by truncating the trained vectors. We thus truncated the trained 16,000-dimensional representations to 8000, 4032, 2048, 1024, 512, 256, 128, and 64 bits and computed performance metrics for each of the resulting vector spaces.

Querying the vector space for familiar concepts presents an opportunity to qualitatively evaluate whether the learned representations are meaningful. Using the Semantic Vectors open source software, the vector space was queried using various drugs, side effects, and compositional concepts to assess whether the space is intuitively interpretable.

Finally, we investigated the proximity of related side effects to each other using a visualization of the side effect vectors, created using Uniform Manifold Approximation and Projection (umap-learn package^14,27^, version 0.3.7), which projects multi-dimensional representations into two dimensions. Individual side effects were colored according to their class as listed in the downloaded Decagon datasets. Many side effects did not belong to any category, and some categories had few or no side effects in them. Of the remaining categories, we visualized the 6 most frequently occurring side effect classes.

Code and data for all preprocessing, training, and evaluation steps are available on Github^28,29^.

## Results

The performance metrics for ESP and Decagon, averaged over 963 side effects, are shown in Table 2. The mean AUROC for the 8-epoch ESP model was 0.903 (range 0.841-0.977); the mean AUPRC was 0.875 (range 0.779-0.976); and the mean AP@50 was 0.865 (range 0.550-1.0). These results exceed both the published Decagon results (in Table 2) and those of our locally-trained Decagon model after 8 epochs: mean AUROC of 0.855 (range 0.259-0.949); a mean AUPRC of 0.793 (range 0.366-0.934); and a mean AP@50 of 0.638 (range 0.066-0.950).

**Table 2.**
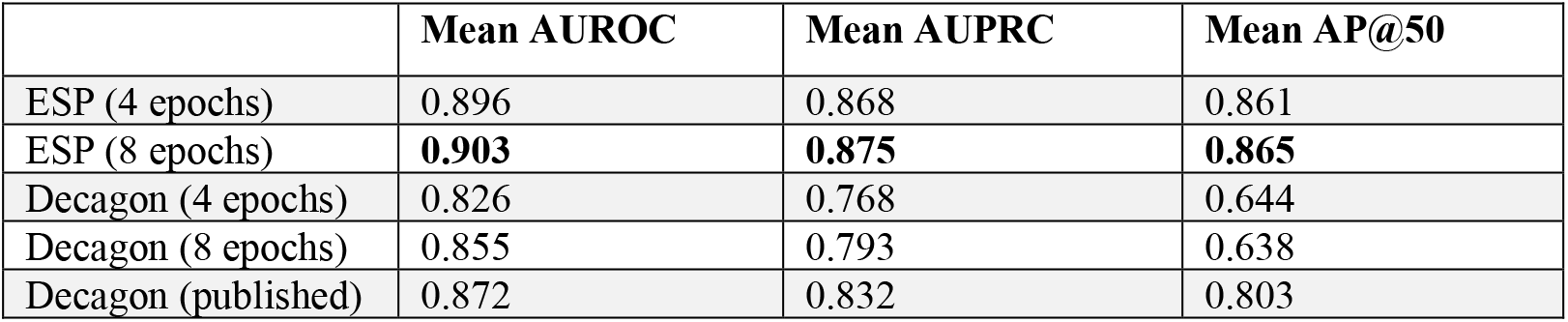
Performance metrics for ESP and Decagon.

Our 4-epoch ESP model took roughly 3.5 hours to train with 16,000 dimensions (bits), including setup time and time spent making predictions over the test set. Each epoch took an average of 50 min. 4 epochs of training Decagon were completed in 6 days and 4 hours including setup and test set prediction, with an average of 36 hours per epoch.

The mean area under the receiver operating characteristic curve over 963 side effects for different vector dimensionalities, created by truncating the 16,000-dimensional vectors, are shown in Figure 2.

**Figure 2.**
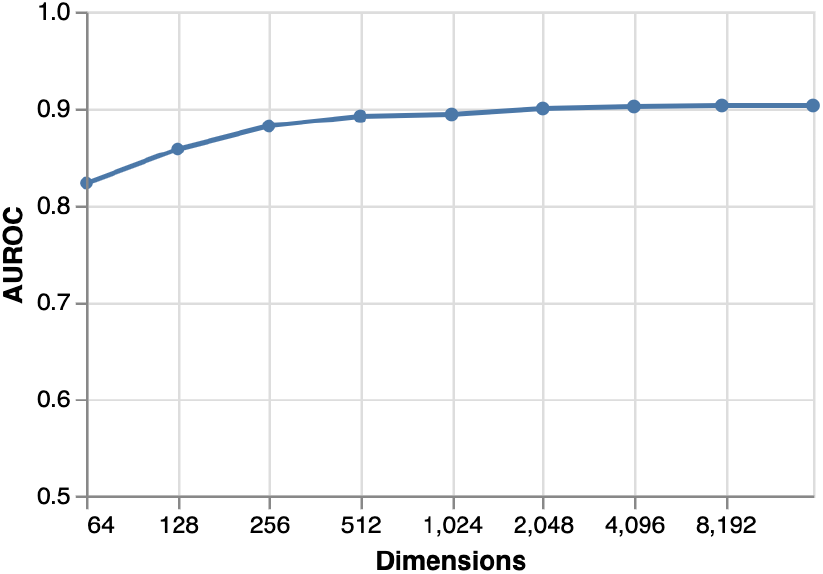
Mean AUROC (over 963 side effects) by vector dimensionality.

Example queries of the trained vector space are shown in Table 3. The search returns the vectors that most similar to the query term, along with a similarity score.

**Table 3.**
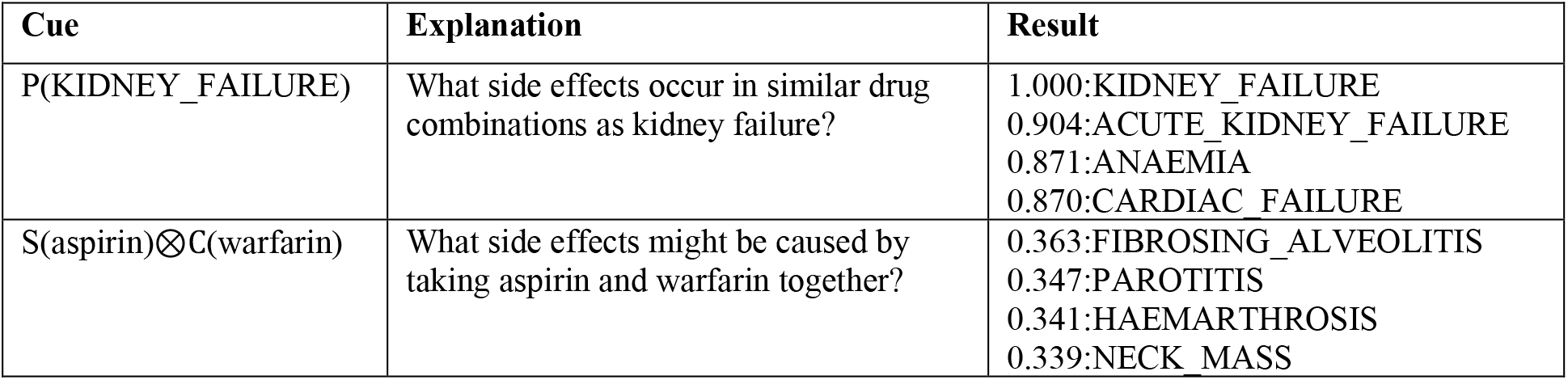
Example searches of the predicate vector space. S,C,P: Semantic, Context and Predicate Vectors

A visualization of how vectors for side effects that are similar to each other are located with respect to one another within the learned embedding space is shown in Figure 3.

**Figure 3.**
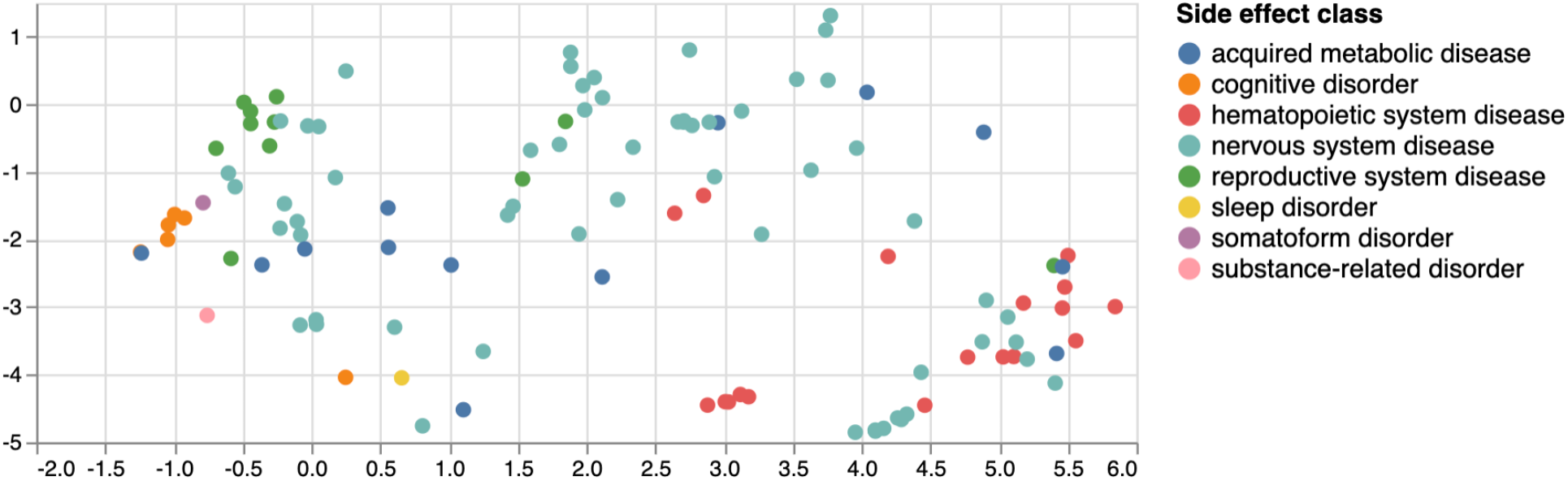
UMAP clustering of a subset of side effect groups. Categories are from Decagon’s published data sets.

## Discussion

ESP outperforms Decagon (both best published performance and our locally-trained model) when predicting polypharmacy side effects by all measures. Furthermore, ESP achieves this performance while training a factor of 43.2 times faster per epoch than Decagon, and for 8 epochs only. The best performing Decagon models were run up to 100 epochs (the number of epochs prior to early stopping was unfortunately not documented). We restricted our locally-trained model to a maximum of 8 epochs on account of resource constraints, which explains the modest difference in performance from previously published results. Additionally, we note that ESP produces re-usable embeddings of the same dimensionality drugs and side effects, which can be used to answer various questions using vector symbolic architecture operations. Using this approach, as well as projection visualization using UMAP, we were able to interpret the embeddings produced by the model.

The high performance that ESP was able to achieve is notable. Considering that Decagon substantially outperformed several baseline approaches, the improved performance of ESP over Decagon puts this model among the top in class. The multimodal network representation and modeling of multiple side effects simultaneously, which allows sharing of model parameters across side effects, is an advantage of Decagon over some alternative polypharmacy side effect prediction models that is shared by ESP, and may be in part responsible for improved performance as the model can learn to generalize across similar concepts (such as ‘kidney failure’ and ‘acute kidney failure’, shown in Table 1).

While training time may not be the most significant factor in determining which models are superior when abundant computational resources are available, a faster model is preferable if degradation of accuracy is not a tradeoff that has to be made. Less extensive hardware requirements enable academic researchers with relatively modest computational resources to reproduce and build upon extant research, thus enabling scientific progress through improved equity. But even for those research groups with high-end computing clusters and multiple GPUs at their disposal, it should be considered that training complex neural network models can incur significant energy costs, with carbon footprints equivalent to as much as five times that of an average car, including manufacture^30^. In comparison, ESP can be trained on commodity laptops or servers. Financial and environmental costs aside, one may argue that resource intensive computations will become progressively easier as processing power increases in accordance with Moore’s law (at least for some time^31^); however, this is a difficult case to make in informatics, where much work is based on data sources that increase in size at an incredible pace themselves^24^. As we harness biomedical literature, electronic health record data, social media sources, and adverse event reports, we also observe their sustained and sometimes explosive (e.g. EHR and patient reported data) growth. Currently, DDI prediction models should be retrained whenever new adverse event report data become available (e.g. weekly or monthly). With the advent of large volumes of patient-level data, DDI prediction models have the potential to be personalized to individual patients’ demographics, health status, and genetic makeup, requiring scores of different models to be trained and re-trained, further increasing training costs. ESP conforms to previously formulated recommendations to prioritize development of efficient models^30^ and can help promote research equity by allowing the use of commodity hardware.

The ability to learn representations that correlate with human judgment of similarity between concepts is an advantage of ESP^16^. In this work, we were able to demonstrate that embeddings learned for related side effects, such as kidney failure and acute kidney failure, are similar (Table 3). However, other types of organ failure are also retrieved by this search. A possible explanation is that patients with multi-organ failure appear in FAERS, and so different sorts of organ failure may become associated with one another. If these organ failures are attached to reports containing the same drug pairs (e.g. drugs used in sepsis), a statistical model will find associations between these drugs and every sort of organ failure. The similarity-based search for side effects using the bound product S(aspirin)⊗C(warfarin) yielded, among others, alveolitis, parotitis, and hemarthrosis. Development of lung disease has previously been reported to be associated with aspirin treatment^32^; additionally, warfarin has been characterized as a drug causing salivary gland swelling and pain^33^. Aspirin’s and warfarin’s possible roles, respectively, in exacerbating these adverse events has yet to be established. We also found similarity of S(aspirin)⊗C(warfarin) with hemarthrosis, a bleeding disorder, which is reasonable, considering that warfarin and aspirin are both anticoagulants. We note that similarity search results may refer to indications rather than side effects. Even though Tatonetti et al.^5^ applied corrections for confounding when creating the TWOSIDES set, with the goal that indications would not appear as side effects, it appears that this effect may not have been entirely eliminated, resulting in our model learning from indication-related statistical patterns remaining in this dataset.

Not all side effect representations for a given side effect category clustered together well in our projection visualization. For example, side effects in the reproductive system, cognitive disorder, and hematopoietic system effect classes are well separated from each other, but nervous system side effects are found throughout the vector space. We suspect that one contributing factor for this is the limited number of types of data in the training set. The only information that ESP has an opportunity to encode into the concept embeddings are polypharmacy side effects and drug target information. As discussed above, side effects appear to cluster because they are actually indications common to particular drug combinations – an additional complicating factor. This was also discussed by Zitnik et al., who reported that many side effects tend to co-occur in this particular data set^11^. As a result, co-location of embeddings in space is not as interpretable as it might be for richer data sets, with more concept and relation types, which might contribute to more meaningful similarities between the various side effects being learned. Previous applications of ESP have involved rich data sets with multiple relationship types and a number of different types of concepts, which resulted in more readily interpretable similarities.

The parsimony of the ESP model is a desirable feature. While Decagon utilizes approximately 87 million trainable parameters, which are each 32-bit floating point numbers – a total of approximately 2.8 billion bits of training parameters – using 16,000-dimensional vectors, ESP performs better with only about 36 million single bit parameters, which is over 77 times fewer bits of representational capacity. In fact, even with 1,024 dimensions, ESP approximates Decagon performance, and this dimensionality corresponds to only 2 million trainable bits of representational power – over *three orders of magnitude* less than Decagon. At 64 bits per embedding – a mere 144,000 bits of trainable parameters – ESP achieves performance that is still higher than the baseline approaches documented by Zitnik et al. This suggests that the vast representational capacity of a trained Decagon model is unnecessary for the task of predicting DDIs that appear in TWOSIDES. More stringent evaluations may be required to reveal any advantages in performance that its larger capacity and more computationally demanding training procedure may confer.

This work has several limitations. Firstly, it is important to note that the data used for training may poorly approximate ground truth. TWOSIDES is a database of information mined from text sources. While a small number of novel drug interactions reported in TWOSIDES have been corroborated through investigation of relevant patient records as well as laboratory experiments^34^, the database contains many unvalidated associations, including some unintuitive side effects that may not withstand curation, such as ‘Mumps’ or ‘Fracture’. These side effects do give us an idea of what the drug interactions may be; for instance, if mumps is increasingly observed in patients taking a certain combination of drugs, it may be plausible to infer the polypharmacy side effect to be a weakening of the immune system. Similarly, if fractures have increased prevalence in the patient population taking particular drugs in combination, we may consider the idea that the interaction of the drugs causes decreased bone density. Yet, using a curated reference set of drug interaction data is preferable over using data obtained through data mining. As discussed above, making models directly comparable is a challenge, which is why we chose to use the same datasets as a comparable previous approach.

Another limitation of this approach is that directionality of relationships between concepts is not encoded. DDIs may be considered symmetric relationships between drugs, as drug A interacting with drug B would also mean that drug B interacts with drug A. Here, we reverse and append all DDI triples in the dataset during preprocessing, creating a separate side effect type representing the “reverse relationship”, as Zitnik et al. did. In other words, for each edge (with a side effect specific edge type) pointing from drug A to drug B, we also add an edge from drug B to drug A, which has a separate, new edge type. While the additional triples (edges) benefit the training of drug embeddings, the predicate embeddings to which they contribute (redundant side effect representations) are not used when making predictions. Conversely, the inverse relationship, despite being known to be true, is not encoded in the retained side effect embedding. Similarly, drug target triples are reversed and appended to the dataset with a new edge type. In the drug target case, this matches the directionality of the relationship: if drug A targets protein B, we cannot also say that protein B targets drug A. Directionality would be obscured if we reversed and appended the drug target triples with the same edge type. Finally, protein-protein interaction (PPI) triples, which are inherently symmetric, are treated yet differently. Decagon quadruples all PPI triples, first by reversing the order and then by creating an alternative relationship, such that the nodes for protein A and protein B are connected by a total of 4 edges: 2 “interacts_with” edges, one pointing from A to B and one from B to A, and 2 “reverse_interacts_with” edges, one pointing from A to B and one pointing from B to A. While ESP can encode directionality directly we did not explore this here and instead chose to follow the same input data preprocessing protocol as Zitnik et al. in order to facilitate direct comparability.

In this research, we have followed the training and evaluation protocol reported by an earlier study as closely as possible for the purpose of comparative evaluation, leveraging both written descriptions of the protocol and publicly available data and software. However, during this process, we found some details in the reported protocol ambiguous, and found apparent inconsistencies between the approach described in the paper and the code it refers to. For instance, the number of side effects and proteins reported by Zitnik et al. does not exactly match the number we found in the data ourselves (963 side effects with 500 or more drug pairs, compared to the reported 964; 19081 unique drug targets compared to the reported 19085). Also, neither cross validation nor early stopping are implemented in the publicly released code, though both methods were reported in the paper. The 10% of data reserved as validation set consequently went completely unused in the published code. As a result of the computational demands of training Decagon, a test of statistical significance, which would require Decagon and ESP to each be trained repeatedly, was not performed. Additionally, we found that all data were split into training and test sets, including protein-protein interaction and drug-target data, even though we are not concerned with making predictions for these data subsets. This equates to discarding a portion of the protein-protein and drug-target data without good reason, other than allowing a simple 80/10/10 split across all types of triples. While these discrepancies were reconciled by our group to our best effort, subtle differences in our evaluation pipelines may not have been totally eliminated.

This project has established a foundation for future work. Having demonstrated the accuracy of ESP as compared to previously reported models while keeping the input data constant, we plan on applying the technique to datasets that are (at least partially) curated rather than statistically derived, e.g. the interaction designations in FAERS reports or the reference set published by Ayvaz et al.^35^. While accuracy measures such as AUROC and AUPRC may not be affected by this change, the resulting model would presumably have more validity when applied to real data.

Incorporating additional knowledge may also present an opportunity to improve this modeling approach. ESP is a method for encoding a knowledge graph; thus, we can generate embeddings for any number of concepts and relationships based on their connections to each other. Many sources of knowledge regarding drugs and their interactions may be relevant. For example, drug classifications, molecular structure information, and additional information about drug target molecules may allow us to train more comprehensive embeddings which could even be applied to new drugs prior to the appearance of any adverse drug event reports. On the patient level, ESP models for predicting DDIs may help make strides in precision medicine, e.g. by incorporating patients’ individual genetic profiles in the model. Training patient specific models would be made feasible in part due to ESP’s efficiency in both training time and model complexity, each being only about 1-2% of what previously reported models require.

## Conclusion

Many individuals are at risk of experiencing adverse events caused by drug-drug interactions. Because DDIs are poorly characterized before drugs are released to market, efficient and accurate analysis of post-market surveillance data is critical to patient safety. ESP, a representation learning method with demonstrated utility in the pharmacovigilance space, provides an alternative approach to DDI prediction. In this work, we have shown that application of ESP to adverse drug event data can yield accurate predictions of polypharmacy side effects, with performance exceeding that of the best reported machine learning models on this task. Being considerably more lightweight in terms of required compute power and model complexity than previously published approaches, ESP can be expected to scale up without imposing undue financial and ethical burdens on researchers, particularly as the volume of data, number of models to train, and frequency of re-training increase. Additionally, we have demonstrated that ESP produces meaningful representations of biomedical concepts that can enable biomedical discovery in novel ways, providing insights that pave the way for further development of ESP models for pharmacovigilance and other applications.

## Acknowledgments

This work is supported by a training fellowship (Grant No. 3T15 LM007442, Biomedical and Health Informatics Training Program) and U.S. National Library of Medicine Grant (R01LM011563), Robust Inference from Observational Data with Distributed Representations of Conceptual Relations.

